# METTL3 inhibition attenuates the macrophage type I interferon response

**DOI:** 10.64898/2026.08.14.744930

**Authors:** Edward M.C. Courvan, Cody J.S. Hecht, Frederick Longshore-Neate, Luisa Macedo de Vasconcelos, Roy Parker

## Abstract

Macrophages coordinate inflammatory responses to changing tissue conditions and invasion by pathogens. Post-transcriptional regulation of the transcriptome via RNA decay is a crucial element of these responses. N^6^-methyladenosine (m^6^A) deposition on mRNA constitutes one such post-transcriptional pathway, where the methyltransferase METTL3 and the oxygen dependent eraser ALKBH5 regulate m^6^A levels. We used TimeLapse-seq to measure transcriptome-wide abundance, decay, and transcription in THP1 macrophages comparing the effects of the METTL3 inhibitor STM2457 and hypoxia. We found no evidence that hypoxia and STM2457 affect RNA decay by a shared mechanism but discovered that METTL3 inhibition transcriptionally downregulates the interferon response through STAT1. Consequently, STM2457 treatment makes both THP1 and peripheral blood macrophages are more permissive to infection by the human coronavirus OC43. We conclude that macrophages depend on METTL3 activity to support the interferon response, an effect that appears distinct to macrophages and suggests METTL3 inhibitors could control macrophage driven inflammation.

## Introduction

Mammalian gene expression relies on post-transcriptional mRNA control, which includes the co-transcriptional deposition of N^6^-methyladenosine (m^6^A) on nascent mRNA by methylase METTL3^1^. Transcripts are regulated downstream by an array of m^6^A “reader” proteins that modulate mRNA translation and stability^2^. Though specific functions of m^6^A appear to be dependent on cell type and biological context, inhibition of m^6^A synthesis by METTL3 has become an attractive therapeutic target for a broad spectrum of cancers^3–6^. Regulated RNA decay plays an important role in macrophage biology, leading us to hypothesize that METTL3 inhibition could have consequences for macrophage function^7^.

The METTL3 inhibitor STM2457 is a first-in-class compound identified as a potential treatment for AML^4^. STM2457 restricts AML proliferation by depleting m^6^A from transcripts that promote cell division and subsequently reducing their translation efficiency^4^. In other tumors, STM2457 treatment translationally downregulates EGFR, pathways that manage oxidative stress, and glycolysis via the IGF2BP proteins^8–10^. STM2457 can also affect tumor suppressor expression by preventing m^6^A-dependent decay via the YTH proteins^11,12^. This potential to regulate oncogenic and tumor suppressive factors explains the broad spectrum of STM2457 treatment possibilities but also raises questions about how METTL3 inhibition may affect other cell types in the tumor environment and elsewhere in the body.

We became intrigued by the idea that m^6^A levels might be regulated by hypoxia, which is a common element of the tumor microenvironment^13^. The m^6^A eraser ALKBH5 has demonstrated dependence on oxygen, implying that hypoxia could slow removal of m^6^A^14^. Hypoxia can lead to accelerated RNA decay in macrophages under inflammatory conditions^15^. Thus, we hypothesized that changing m^6^A levels could drive changes in RNA decay rates for macrophages in the tumor microenvironment.

Here, we use TimeLapse-seq to measure decay rates in THP1 macrophages treated with STM2457, hypoxia, or both together. We arrive at the conclusion that METTL3 inhibition and hypoxia affect RNA decay rates by separate mechanisms in macrophages. Unexpectedly, we discover that METTL3 inhibition with STM2457 reduces STAT1 mediated transcription. Ultimately, we find that METTL3 inhibition suppresses the interferon response and promotes OC43 viral proliferation in macrophages, a finding that appears to be unique to this cell lineage.

## Results

### METTL3 inhibition and hypoxia both downregulate interferon related genes

We began by deploying TimeLapse-seq to systematically measure abundance, decay, and transcription changes in THP1 macrophages, where we compared the effects of STM2457 treatment, hypoxia, and both STM2457 and hypoxia^16–18^. Briefly, TimeLapse-seq uses 4-thiouridine (s^4^U) metabolic labeling to mark newly transcribed RNA at the end of a time course. s^4^U incorporated into RNA is then chemically modified before library preparation, where these bases are read as mutated to cytosine rather than uridine by the reverse transcriptase^16,18^.

First, we inspected the effects of STM2457 on transcript abundance (Figure 1A&B). Our expectation was that METTL3 inhibition would lead to gene expression changes driven by downregulation of the same proliferative transcription factors impacted when STM2457 is used on cancers^4,5,8,9,12^. Instead, we were surprised to find that highly upregulated transcripts are dominated by poorly annotated long non-coding RNAs (lncRNA; Figure 1A). Examining genome browser tracks indicated that many of these lncRNAs and other transcripts are upregulated as a result of readthrough transcription from upstream loci (Figure S1A&B).

**Figure 1.**
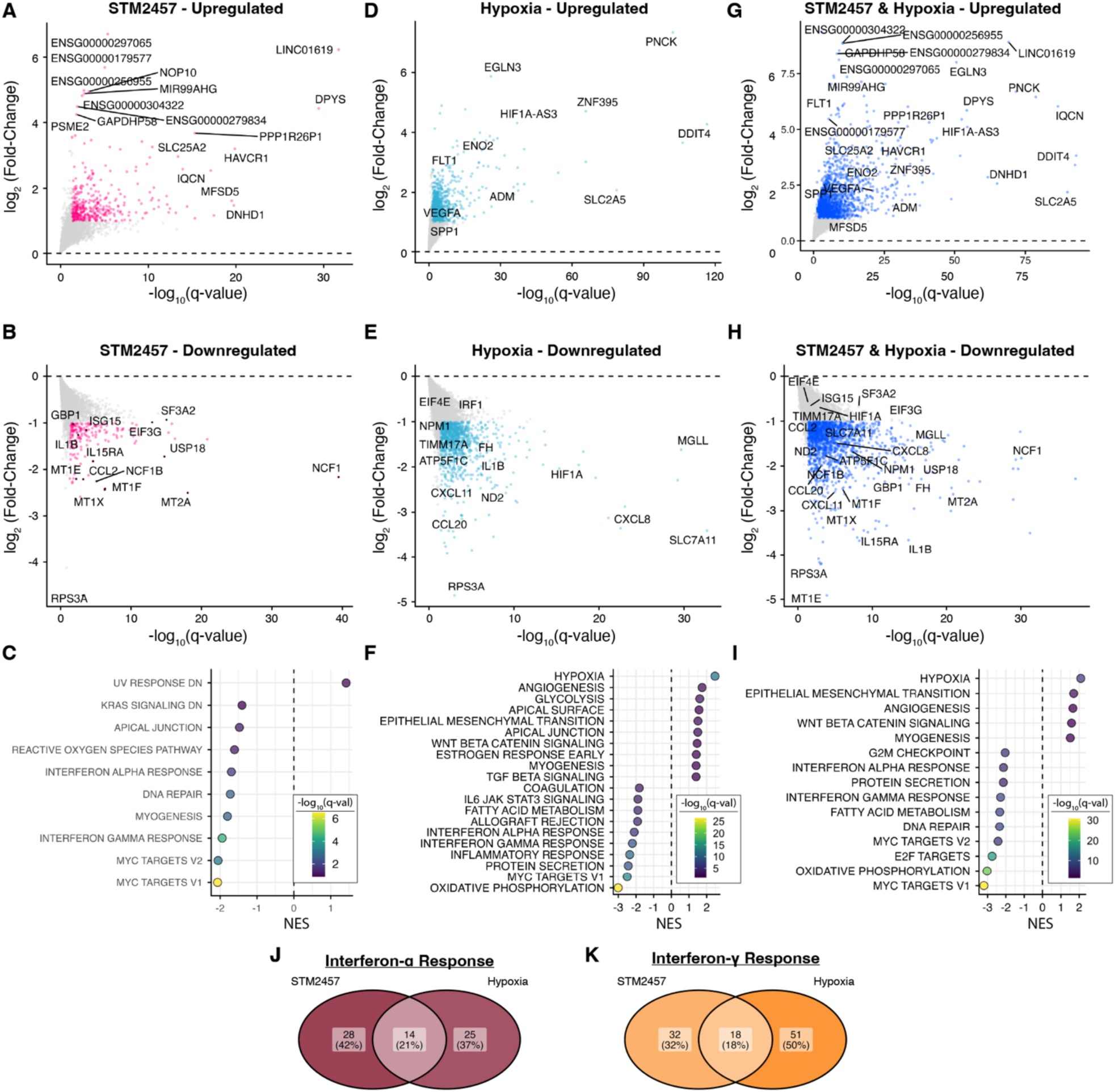
STM2457 and hypoxia regulate transcript abundance independently. Differential expression analysis on THP1 macrophages treated with STM2457 (A-C), 1% sea-level equivalent O_2_ hypoxia (D-F), or both simultaneously (G-I). Upregulated transcripts (A,D,G) are shown in color if a transcript is upregulated by more than two-fold. Downregulated transcripts (B,E,H) are shown in color if downregulated by more than two-fold. Both apply a q-value threshold of 0.05. GSEA was performed on gene lists ranked by differential expression (C,F,I), and the top and bottom ten gene sets from the MSigDB hallmark gene set collection were filtered by a 25% false discovery rate threshold and plotted by normalized enrichment score (NES). J&K) Venn diagrams of leading-edge genes for the interferon-α response (J) or interferon-γ response (K).

Transcripts downregulated by METTL3 inhibition include macrophage inflammation markers like metallothioneins and components of the phagocyte NADPH oxidase complex (Figure 1B). We found this surprising, as other data have shown m^6^A depletion to promote inflammatory signaling via increased dsRNA sensing^19,20^.

Using gene set enrichment analysis (GSEA) with the MSigDB hallmark collection of gene sets, we found that both interferon-α and γ related genes were enriched among downregulated genes alongside MYC targets and reactive oxygen pathways (Figure 1C). Only the UV response was identified as upregulated (Figure 1C)^21,22^.

Hypoxia alone also induces widespread changes to gene expression. Induced genes include: EGLN3, which encodes the prolyl hydroxylase that mediates HIF1A protein degradation; and HIF1A antisense transcripts, both indicators of a normal hypoxic response (Figure 1D). Downregulated transcripts include: HIF1A itself; transcripts related to oxidative phosphorylation machinery; and inflammatory cytokines such as IL1B, CXCL11, and CXCL8 (Figure 1E).

GSEA shows that interferon response gene sets are also downregulated by hypoxia (Figure 1F). As expected, hypoxia, angiogenesis, and glycolysis gene sets are upregulated, whereas metabolic pathways such as fatty acid processing and oxidative phosphorylation gene sets are downregulated in addition to both interferon gene sets.

In combination, METTL3 inhibition by STM2457 and hypoxia reproduce the gene expression changes of each individual stimulus simultaneously (Figure 1G&H). By correlating changes in decay rates in STM2457 and hypoxia to published m^6^A densities as well as comparing differential expression, we judge METTL3 inhibition and hypoxia to be mechanistically orthogonal (Figure S1C-F). GSEA reveals that most of the pathways regulated by METTL3 inhibition and hypoxia alone are directionally matched by both together (Figure 1I). Notably, both METTL3 inhibition and hypoxia downregulate interferon response gene sets. We compared the subsets of genes driving the interferon-α and γ response gene sets and found little overlap between genes affected by either STM2457 or hypoxia (21% and 18% respectively; Figure 1J&K).

We conclude that METTL3 inhibition by STM2457 downregulates both type I and type II interferon response related genes at the transcript level. Hypoxia also downregulates interferon response genes, but this effect arises through a separate module.

### METTL3 inhibition blocks interferon related gene expression by a transcriptional mechanism

We next asked whether the kinetic data produced by TimeLapse-seq could provide insight into the mechanism of interferon related gene downregulation. For STM2457 treatment alone, we observe that most transcripts experienced decreased decay, with the notable inclusion of the YTHDF2 transcript itself, which can be autoregulated by m^6^A levels (Figure 2A)^23^. From our previous work, we expected hypoxia to destabilize the transcriptome, which we indeed see (Figure 2B)^15^. The combination of STM2457 treatment with hypoxia leads to a centering of global turnover rates, with many of the individual transcripts maintaining their relative change in stability (Figure 2C). Viewed together, we see that STM2457 and hypoxia have opposing effects which moderate one another (Figure 2D).

**Figure 2.**
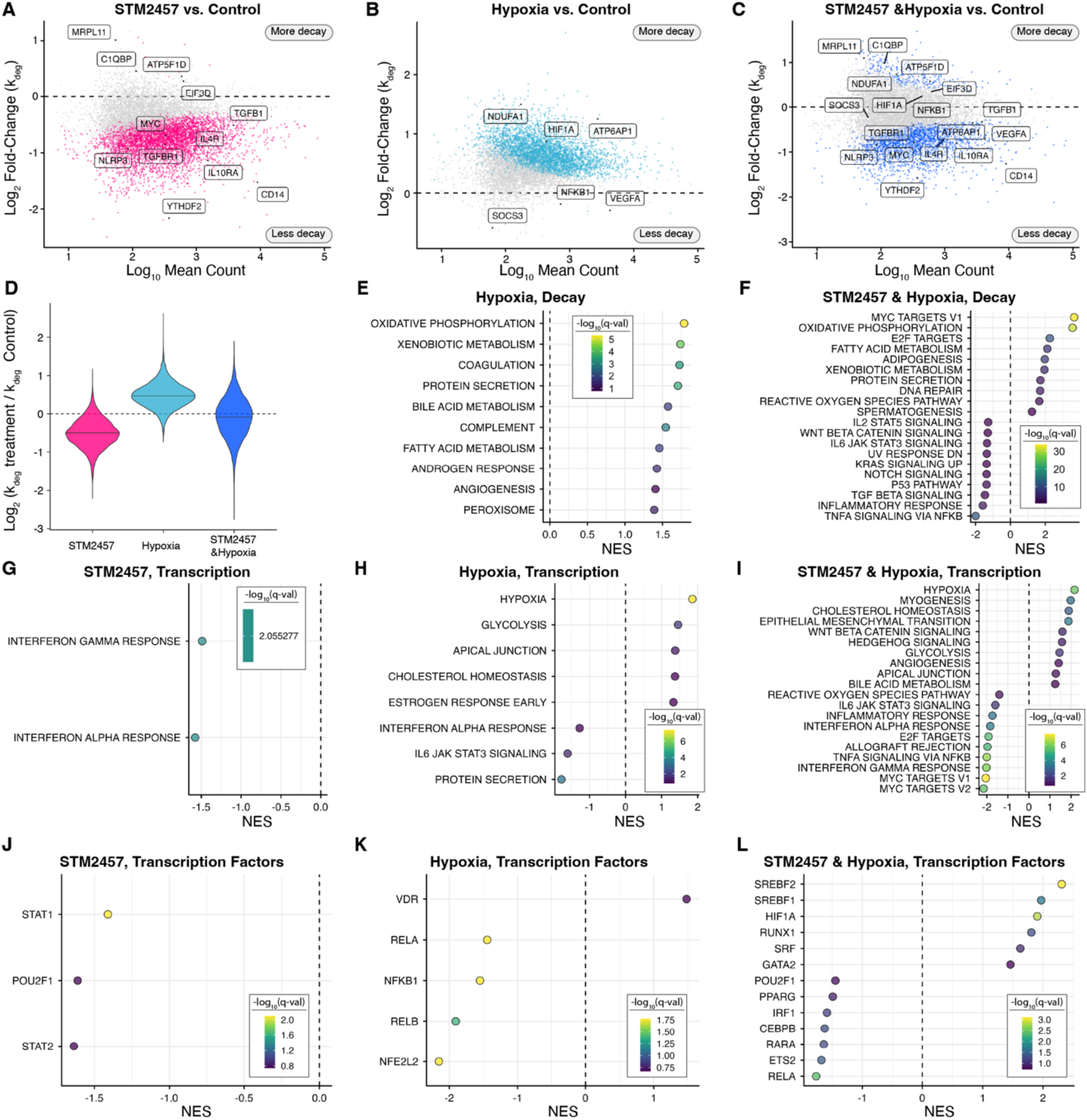
METTL3 inhibition leads to transcriptional repression of interferon stimulated genes. A-C) Differential decay vs. abundance for each condition measured with TimeLapse-seq where A) shows METTL3 inhibition with STM2457, B) shows hypoxia, and C) shows both treatments together. Each point represents a transcript, where points in color meet a threshold of q < 0.05. D) Summary of differential decay derived from TimeLapse-seq; solid line indicates the median of the distribution. E&F) GSEA of differential decay plotted by normalized enrichment score (NES) for E) hypoxia and F) STM2457 and hypoxia. GSEA of STM2457 treatment alone produced no gene sets meeting the false discovery rate (FDR) threshold of 25%. Gene sets taken from the MSigDB hallmark collection. G-I) GSEA, again using the hallmark collection, of differential transcription derived from differential expression and differential decay data for STM2457 (G), hypoxia (H), and both together (I). J-L) GSEA on differential transcription using the DoRothEA database of transcriptional activators. All GSEA uses a cutoff of 25% FDR and displays up to ten pathways or transcription factors.

We returned to GSEA to ask whether there is any pathway specificity embedded in transcripts regulated by differential decay. We were surprised to find that no gene set enriched in our STM2457 dataset, suggesting that METTL3 inhibition agnostically stabilizes mRNAs with respect to pathway (Figure S2A). In contrast, hypoxia led to enrichment of several pathways, the foremost being oxidative phosphorylation (Figure 2E&S2B). In combination, STM2457 and hypoxia differential decay enrich many of the same pathways as hypoxia alone, with the addition of MYC targets and TNFα signaling gene sets as more susceptible to decay and stabilization, respectively (Figure 2F&S2C). From this, we interpret that the stabilization effects of METTL3 inhibition are largely nonspecific. The pathway regulation embedded in hypoxic destabilization confirms global changes in RNA decay regulation can carry specificity, but this is unlikely to be the case for m^6^A mediated decay.

We were struck by the fact that neither interferon gene set is present among enriched gene sets in any condition. Thus, we ruled out RNA decay as a driver of reduced interferon related gene expression in either STM2457 treatment or hypoxia.

Given that most pathways affected by METTL3 inhibition or hypoxia are seen as affected by the combination of the two, we strongly suspected that METTL3 inhibition and hypoxia affect decay by independent mechanisms. When we compared published m^6^A levels measured by GLORI (<u>g</u>lyoxal and nitrite-mediated deamination of unmethylated adenosines), either in hypoxia or normoxia, to our differential decay datasets, we find that only METTL3 inhibition induced changes in decay correlate to m^6^A levels (Figure S2D-F). Hypoxia induced changes in decay are not correlated to m^6^A modification rates, leading us to conclude that METTL3 inhibition and hypoxia regulate RNA decay by independent mechanisms.

We found that the downregulation of interferon response genes seen in our total RNA-seq analysis for STM2457 is explained by transcriptional downregulation (Figure 2G). This analysis is possible because TimeLapse-seq characterizes differential expression and decay, allowing us to infer differential synthesis (Δabundance = Δsynthesis - Δdecay; assuming steady state dynamics)^18^. Using GSEA, we asked which pathways were dominated by transcriptional regulation. STM2457-induced transcriptional regulation exclusively depletes the interferon response gene sets. Likewise, interferon-α response genes downregulated in hypoxia can also be assigned to a transcriptional mechanism, and these results are borne out in the combined treatment (Figure 2H&I).

We compared our differential synthesis results to transcription factor annotations in the DoRothEA collection of transcriptional regulons to determine what transcriptional modules are downregulated by either METTL3 inhibition or hypoxia^24^. Here, we were intrigued to find that differentially transcribed genes after METTL3 inhibition are predominantly STAT1 and STAT2 targets (Figure 2J). This was not shared by either hypoxia or the combined treatment (Figure 2K&L). Decreased STAT1 and STAT2 driven transcription strongly suggested that METTL3 inhibition suppresses the interferon response far upstream at the level of the interferon receptors, kinases and the STAT transcription factors themselves.

### METTL3 inhibition decreases expression of the type I interferon receptor module

To confirm that METTL3 inhibition affects the interferon response, we performed a second RNA-seq experiment comparing STM2457 treatment to control in both resting and interferon-α stimulated THP1 macrophages. This experiment recapitulated the downregulation of both interferon-α and interferon-γ response genes in response to STM2457 treatment (Figure 3A&B; Figure S3A&B). When we compare STM2457 to control in the context of 1 ng/mL interferon-α for 4 hours, the suppression of interferon related genes is more pronounced (Figure 3B&C). Thus, we conclude that METTL3 inhibition suppresses the macrophage interferon response.

**Figure 3.**
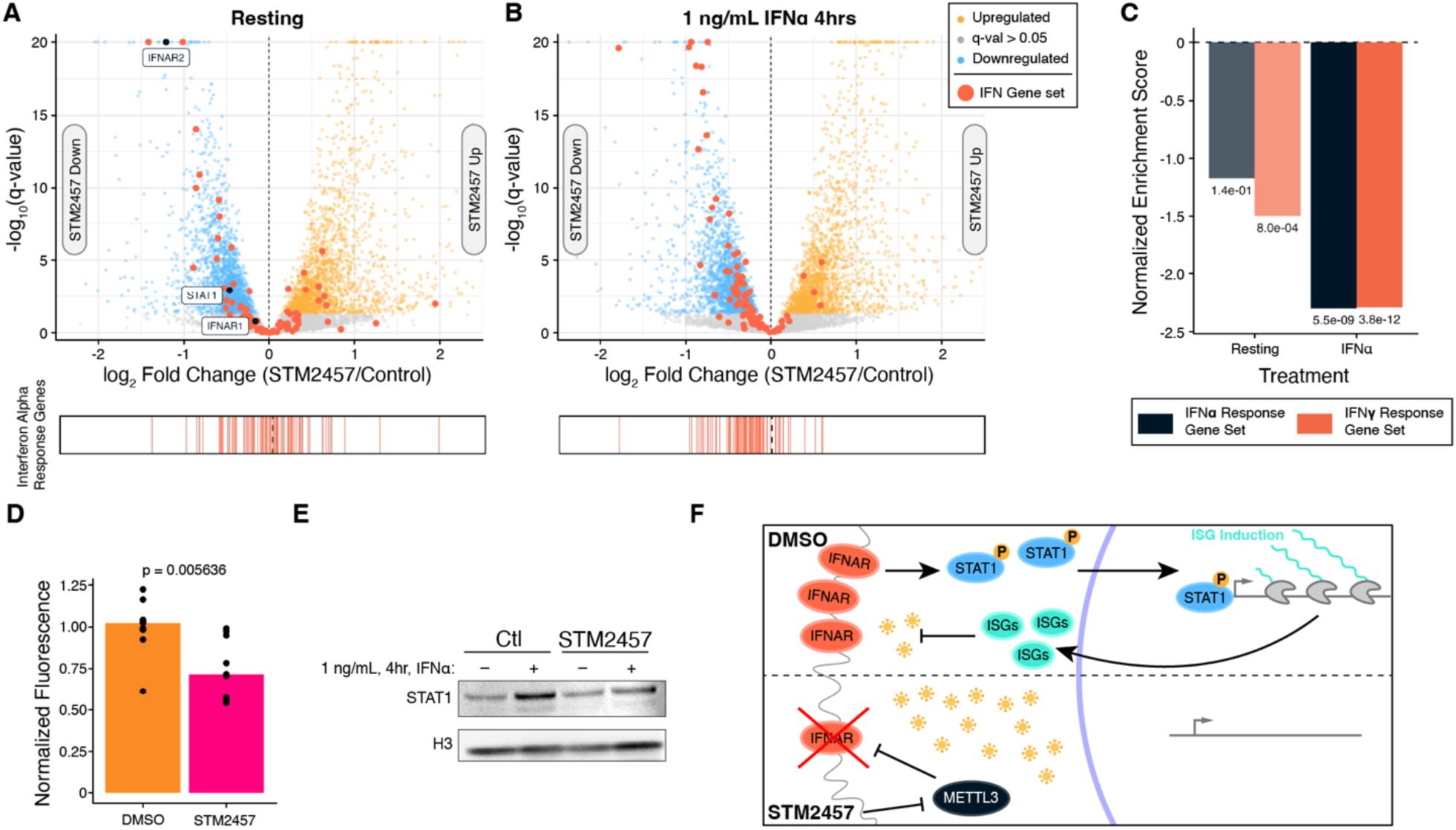
METTL3 inhibition by STM2457 inhibits the interferon-α response at the receptor level. A&B) PolyA RNA-seq of resting THP1 macrophages (A) or THP1 macrophages treated with 1 ng/mL interferon-α for four hours (B). Significance determined by an FDR threshold of 25%. Transcripts belonging to the MSigDB hallmark “Interferon Alpha Response” gene set are noted in red and scored in the strip plot below. C) GSEA-derived normalized enrichment scores for resting and interferon-α treatment for RNA-seq datasets in A&B. D) Flow cytometry for interferon alpha/beta receptor 2 in THP1 macrophages in either control or STM2457 treatment. Reported p value taken from Mann-Whitney U. Fluorescence values were background subtracted and normalized to DMSO control. E) Western blot of STAT1 for STM2457 treatment in the presence and absence of interferon-α. F) Schematic of how STM2457 treatment inhibits METTL3 and subsequently blocks IFNAR2 expression leading to decreased interferon response and a permissive cellular environment for viral replication.

Prior to interferon-α treatment, transcripts encoding IFNAR2, a component of the type I interferon receptor, and STAT1, the dominant interferon transcription factor, are downregulated by METTL3 inhibition (Figure 3A). We measured the surface expression of IFNAR2 and IFNGR2 by flow cytometry and found that STM2457 reduces expression of IFNAR2 by 25%, though it does not appear to affect surface IFNGR2 (Figure 3D&S3C). Measuring total STAT1 expression by western blot after interferon stimulation shows that protein induction is impaired by METTL3 inhibition (Figure 3E). From these results, we predicted that METTL3 inhibition would impair the macrophage antiviral response (Figure 3F).

### METTL3 inhibition prevents interferon from priming the macrophage interferon response

To determine how METTL3 inhibition affects the antiviral response in macrophages, we infected THP1 macrophages with the human coronavirus OC43 in four conditions: control, interferon-α primed, METTL3 inhibition with STM2457, and both primed and METTL3 inhibited. We primed for 24 hours to allow full expression of the interferon response and then infected with OC43 for 8 hours before collecting samples (Figure 4A). We measured induction of the interferon stimulated gene ISG15 and found, at this time point, interferon-α induces ISG15 regardless of whether METTL3 is inhibited. OC43 infection alone also induces ISG15, and together, we interpret that THP1 macrophages still mount a response to both interferon-α and viral infection as judged by ISG15 alone, but the response is likely to be muted at the transcriptome level (Figure 4B).

**Figure 4.**
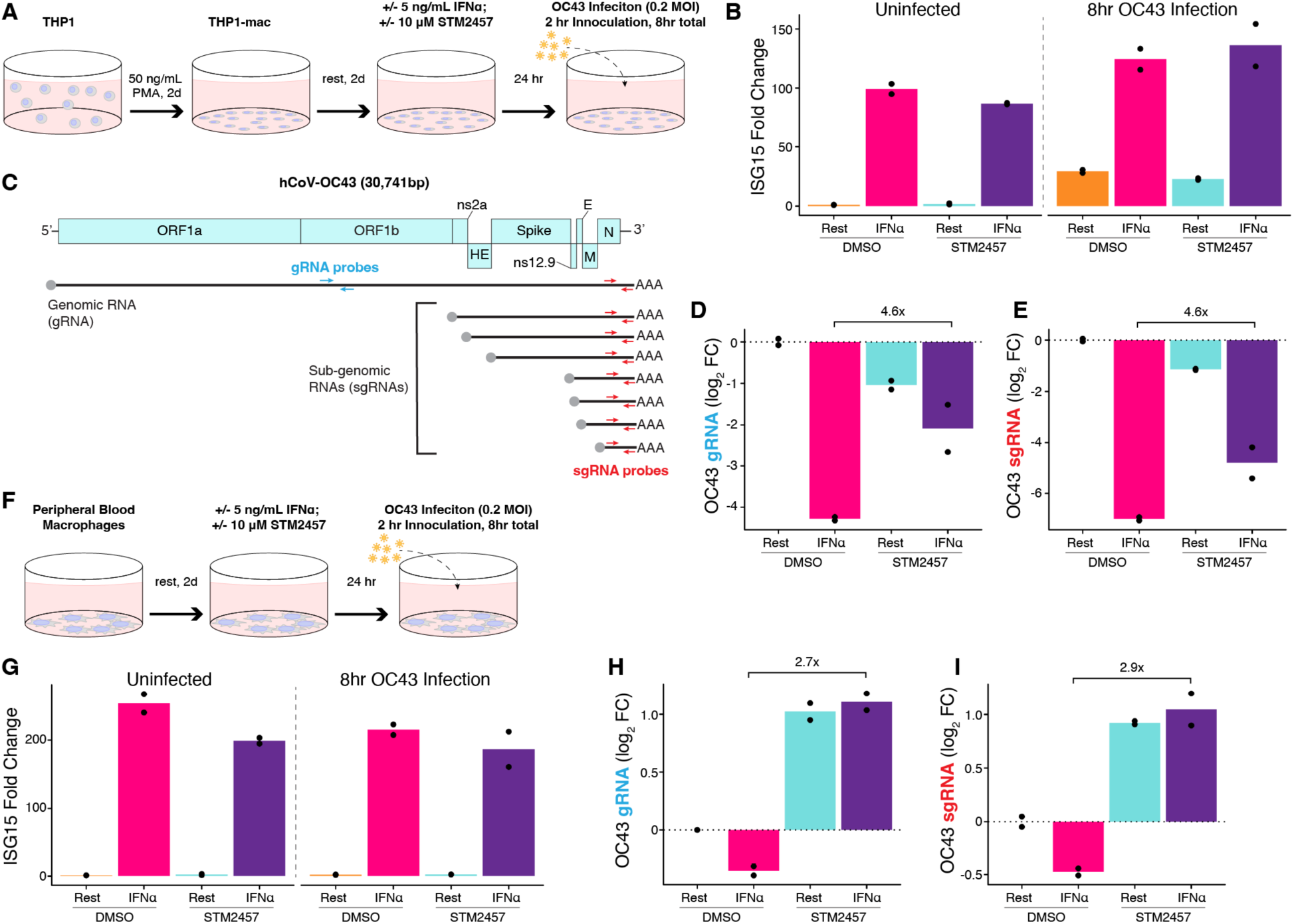
Macrophages are more susceptible to OC43 infection during METTL3 inhibition. A) Schematic of THP1 differentiation with phorbol 12-myristate 13-acetate (PMA), STM2457 and interferon (IFN) treatment, 2-hour inoculation with OC43 and 8 hrs total incubation. B) RT-qPCR for ISG15 across all four treatment conditions in uninfected and OC43 infected cells. C) RT-qPCR strategy for OC43 detection including probe location for genomic RNA (blue; gRNA) only and for the combined genomic and sub-genomic RNAs produced during active infection (red; sgRNA). D) RT-qPCR for OC43 gRNA. E) RT-qPCR for OC43 sgRNA. F) Schematic of treatment and infection of peripheral blood macrophages as in (A). G-I) RT-qPCR as in B, D & E on OC43 infected peripheral blood macrophages. All RT-qPCR values were expression normalized to the 18S ribosomal RNA and plotted relative to the resting DMSO condition.

We assessed viral proliferation via RT-qPCR for both the OC43 genomic RNA and sub-genomic RNAs produced by active infection (Figure 4C). Consistent with interferon’s antiviral effects, we observe that interferon-α priming reduces viral proliferation (Figure 4D&E)^25^. STM2457 treatment alone modestly reduces proliferation, consistent with prior results showing m^6^A promotes coronavirus replication^26^.

We were surprised to find that METTL3 inhibition by STM2457 leads to an impaired interferon priming effect whereby viral loads are approximately 4.6-fold higher than with interferon priming alone (Figure 4D&E). We found the same pattern of effects in a second fully independent replicate (Figure S4A-C).

Finally, we asked whether our results transferred from THP1-derived macrophages to peripheral blood macrophages (Figure 4F). Again, we found that ISG15 was induced by interferon-α with and without STM2457, though we noticed the response was slightly weaker under METTL3 inhibition (Figure 4G). Viral proliferation is more restricted in peripheral blood macrophages than in THP1 macrophages (Figure S4D&E). Peripheral blood macrophages inhibit viral proliferation if primed with interferon-α, though the effect is less than in THP1 macrophages, possibly due to a higher baseline antiviral capacity (Figure 4H&I).

Unlike THP1 macrophages, we found that METTL3 inhibition permits a 2-fold increase in viral proliferation regardless of priming, with that effect rising to nearly three-fold when comparing interferon-α primed macrophages (Figure 4H&I). We arrive at the conclusion that METTL3 inhibition impairs the macrophage antiviral response in both THP1 and peripheral blood macrophages.

## Discussion

Here we have investigated possible interplay between how the METTL3 inhibitor STM2457 and hypoxia regulate the transcriptome. Though both induce global changes in RNA decay in macrophage cell lines, our TimeLapse-seq data is more consistent with these changes being driven by independent mechanisms. Nevertheless, we noted that transcriptional regulation after METTL3 inhibition in THP1 macrophages decreases interferon related gene expression. Ultimately, we find that METTL3 inhibition reduces the induction of type I interferon related genes after stimulation with interferon-α, leading to a reduced antiviral response in both THP1 macrophages and peripheral blood macrophages.

Macrophages appear to be unique compared to other cell types where METTL3 disruption has been studied. Deletion of other members in the METTL3 holocomplex in cardiomyocytes leads to increased interferon signaling via stabilization of IFN transcripts^27^. STM2457 and its newer derivatives increase interferon signaling via an accumulation of dsRNA in cancer cells^19,20^. In contrast, our work shows that macrophages respond to METTL3 inhibition by reducing their sensitivity to interferon. Other work has proposed that m^6^A regulates the stability of KDM6B transcript, a histone methyltransferase known to be involved in macrophage activation, and that STAT1 transcripts themselves are subject to m^6^A-mediated stabilization during macrophage activation^28,29^. Our datasets do not point to these specific mechanisms, but in sum, these studies and ours support macrophages as having different rules than other cell types.

Many biological phenomena have evolved to take advantage of global shifts in a fundamental process. The expression of ATF4, the canonical stress response transcription factor, is a paradigmatic example: global translation shutdown allows the ribosome to bypass the upstream open reading frame that typically prevents expression of the transcript^30,31^. We were surprised to find that IFNAR2 and STAT1 are downregulated by METTL3 inhibition despite a tidal shift toward transcript stabilization. The transcript for IFNAR1, the partner of IFNAR2, has been reported to be m^6^A sensitive in other contexts, but this effect is absent in our data^32^. m^6^A can promote expression through reader proteins that recruit the ribosome to modified transcripts. We find it plausible that in macrophages, the transcripts encoding the type I interferon receptor/kinase/transcription module is dependent on reader proteins like YTHDC2 to promote translation^33^. Whether suppressed interferon related gene expression represents a specific downregulation of the pathway or a broader reprogramming of macrophage phenotype remains to be seen.

STM2457 and its derivatives are being explored for treatment of a variety of cancers, but data on their uses to modulate immune function are more limited^4,5,8,9,11,12^. m^6^A depletion has been shown to reduce T cell mediated allograft rejection^34^. METTL3 inhibitors have been shown to have specific antiviral effects in other cell lines^35,36^. Given that m^6^A depletion with METTL3 inhibitors has effects that are specific to macrophages, we presume that these compounds may have applications beyond cancer treatment. We also note that it may be worth considering macrophage inflammatory responses as a source of potential off-target effects in treatment schemes involving these drugs.

To conclude, we have shown that the METTL3 inhibitor STM2457 results in a reduced response to interferon stimulation in THP1-derived macrophages. We propose that METTL3 inhibition may be an underappreciated approach for regulating innate immune reactivity.

## Acknowledgements

This work was supported by the Howard Hughes Medical Institute, the Damon Runyon Cancer Research Foundation, National Institutes of Health awards T32GM145437 and T32GM142607, and equipment grant S10OD021601. The authors would like to thank all members of the Parker lab for providing feedback and assistance throughout the project.

## Data and Code Availability

All sequencing data is available from GEO under accession GSE343705. All code used for processing TimeLapse-seq and RNA-seq is available at https://github.com/ParkerLabCU/METTL3inhib_Macs_Courvan2026.

## Author Contributions

E.M.C.C. and R.P. conceptualized the study. E.M.C.C. led the investigation, experimentation, and data analysis. C.J.S.H., F.L.N., and L.M.V. assisted in the conducting of experiments. E.M.C.C. wrote the manuscript and all authors contributed to editing and revisions.

## Experimental Methods

### Cell culture

THP1 cells were obtained from ATCC (TIB-202) and maintained in RPMI supplemented with 10% FBS at 37°C and 5% CO_2_. THP1 macrophages were differentiated by plating in complete media supplemented with 50 ng/mL phorbol 12-myristate 13-acetate (PMA; Sigma-Aldrich #P8139) for 48 hours. PMA was washed out and cells were allowed to rest for an additional 48 hours after PMA treatment. Peripheral blood macrophages were obtained from STEMCELL Technologies (#70042) and cultured according to supplier protocols in ImmunoCult-SF Macrophage Medium (#10961). Hypoxic conditions were induced for 24 hours by transferring cells to an incubator (Oxford Optronix Hypoxylab) maintaining 7 mmHg O_2_ (equivalent to 1% O_2_ at sea level) with 5% CO_2_ and balanced with N_2_. m^6^A was depleted by treating cells with 10 μM STM2457 in DMSO (Sigma-Aldrich #SML3360) for 24 hours whereas control cells were treated with only 0.1% DMSO. Interferon-α (Thermo Fisher #300-02BC) treatment was applied at 5 ng/mL for 24 hours (antiviral priming) or 1 ng/mL for 4 hours (all other experiments).

### TimeLapse-seq

Triplicate TimeLapse-seq samples were fed with 500 μM s^4^U (Sigma #T4509) for two hours at the end of the 24 hr time course. One additional sample for each condition was not fed s^4^U to control for mutations across comparable expression levels. RNA was harvested by lifting cells into ice cold PBS and dissociating in Trizol followed by precipitation with isopropanol supplemented with 1 mM dithiothreitol (DTT; Thermo Fisher). Carry over genomic DNA was removed by 1 hour TURBO DNase (Thermo Fisher #AM2239) treatment followed by RNAClean XP cleanup (Beckman Coulter #A63987). RNA was treated with 100 mM sodium acetate, 4 mM EDTA, 5% v/v 2,2,2-trifluoroethylamine, and 10 mM sodium periodate at 45°C to oxidize s^4^U and then purified using RNAClean beads. Each sample was then reduced with 1 mM DTT, 10 mM Tris pH 7.0, 1 mM EDTA, and 100 mM NaCl for 30 min at 37°C followed by a final bead purification before library preparation.

### Library preparation and sequencing

TimeLapse-seq libraries were prepared using the Watchmaker RNA Library Kit with Polaris Depletion according to the manufacturer protocol. Libraries were prepared for deduplication compatibility using the Twist Biosciences UMI Adapter system (Twist #105041). The quality of libraries was assessed by Agilent Bioanalyzer and qPCR. Illumina paired-end 150 sequencing was performed by Azenta’s Genewiz sequencing services and produced a minimum of 22 million reads per sample.

Validation RNA-seq was performed by isolating RNA from THP1 macrophages after treatment with STM2457 and stimulation with interferon-α using the RNeasy purification kit according to manufacturer instructions. Residual DNA was removed with TURBO DNase and RNA was bead purified into Tris-EDTA buffer supplemented with SEQguard Dino Preserve (Plasmidsaurus #PS-SDP-024). Library preparation and sequencing were carried out by Plasmidsaurus and sequenced to between 7.6 – 20 million reads per sample.

### Flow cytometry

Flow cytometry was performed at the University of Colorado BioFrontiers Institute Flow Cytometry core (RRID:SCR_019309) using an Accuri C6 Cytometer using the 488 nm laser with a 533/30 filter set and the 640 nm laser with a 675/25 filter set. THP1 macrophages were harvested by scraping and fixed with 4% paraformaldehyde in PBS (Santa Cruz Biotechnology #sc-281692). Cells were blocked in 3% BSA (Sigma-Aldrich #A9647) supplemented with 0.5% normal goat serum (Abcam #ab7481). IFNAR2 was stained with an Alexa 488 conjugated antibody (Cell Signaling Technology #28668S) and IFNGR2 (R&D Systems #FAB773A) was stained with an APC conjugated antibody in blocking buffer for 1 hour. Cells were washed in PBS three times and then filtered for analysis. 20,000 events were collected per sample. FSC-A and SSC-A gating for intact cells was determined by selecting the dominant population in an unstained control (Figure S5). No multiplet population could be detected by gating on FSC-A/FSC-H parameters. Cell populations for each sample were exported as CSV prior to analysis in R. Fluorescence intensities were background subtracted using single stained control cells and normalized to the mean control value. Significance was assessed using Mann-Whitney U test.

### Western blotting

Western blotting for STAT1 and histone H3 were performed by collecting total protein from treated THP1 macrophages in RIPA buffer and separation by SDS-PAGE. Protein was transferred to nitrocellulose via semi-dry transfer and blocked in 5% milk. STAT1 primary staining (Cell Signaling Technology #14994) was performed overnight, membranes were washed three times in TBST and then stained with HRP conjugated anti-rabbit IgG (Cell Signaling Technology #7074T) before three final TBST washes and imaging. Histone H3 was detected with a HRP conjugated primary antibody (Cell Signaling Technology #12648S) followed by washing and imaging. Representative image shown from one of three biological replicates.

### OC43 infection

OC43 growth and titrations were performed in VeroE6 cells as described previously^37,38^. OC43 infections of PMA-differentiated THP1 macrophages and peripheral blood macrophages were performed in technical duplicate. After priming and/or STM2457 treatment for 24hr, cell media was removed and an OC43 inoculant of 8 × 10^4^ TCID_50_ (MOI 0.2) in complete media containing DMSO, 5 ng/mL interferon-α, 10 μM STM2457, or both interferon-α and STM2457 was added. After 2 hours of viral adsorption at 33 °C, the inoculant was removed, and cells were washed once with complete media. Pre-warmed complete media containing DMSO, 5 ng/mL interferon-α, 10 μM STM2457, or both interferon-α and STM2457 was added to cells for an additional 6 hours at 37 °C. After 8 hours of total infection, cells were washed gently once with PBS, lysed in Trizol (Thermo Fisher #15596026), and frozen at -80 °C until RNA extraction. RNA was extracted with Direct-zol RNA Miniprep (Zymo #R2050) according to the manufacturer’s instructions, including on-column DNase I digestion. RT-qPCR was performed with iTaq Universal One-Step RT-qPCR Kit (Bio-Rad #1725150) in 10 μL reactions containing 2 ng of RNA and 200 nM of forward and reverse primers. Thermocycler settings are as follows: 50 °C for 10 minutes, 95 °C for 5 minutes and 39 cycles of 95 °C for 10 seconds and 60 °C for 30 seconds. A melt curve of 0.5 °C/sec from 65 °C to 95 °C was generated with no off-target amplification observed for any of the primer sets. Following qPCR, relative gene abundance was determined using the 2^-ΔΔCt^ method with 18S as the internal control. After 18S normalization, gene abundance for each sample was compared to Uninfected cells + DMSO (for ISG15) or Infected cells + DMSO (for OC43 viral genes). Primer sets listed in Table S1.

## Quantification and Data Analysis

### Processing raw TimeLapse-seq data

Raw sequencing data was processed using established pipelines (fastq2EZbakR) for processing nucleoside recoding data^16–18^. Briefly, sequencing reads are filtered for PCR duplicates, trimmed of adaptors and aligned using HISAT-3N. We aligned our library to version 49.0 of the GENCODE GRCh38 genome annotation. The resulting output includes expression count files, mutation count files, and genome browser tracks.

### Processing validation RNA-seq data

Plasmidsaurus provides 3’ end counting data via oligo-dT priming library preparation and automated alignment (STAR v2.7), deduplication (UMICollapse v1.1.0), and read counting (featureCounts v2.1.1). The resulting count files were filtered for mRNA counts only.

### Differential Expression Analysis

All differential expression analysis was performed by processing count data through DESeq2^39^. Before running DESeq2, we filtered out any transcript with fewer than 100 counts summed across all samples in each analysis. For TimeLapse-seq data, we first included both s^4^U fed and unfed samples to verify that s^4^U does not drastically alter expression and then proceeded to run comparisons on s^4^U triplicates only. We applied a significance threshold of 0.05 to Benjamini-Hochberg q-values across all datasets.

### Differential decay and transcription analysis

Using the s^4^U induced mutation count matrices, we used BakR^17^ to infer differential degradation for all transcripts with at least 50 reads per sample, comparing STM2457, hypoxia, and combined STM2457 and hypoxia to untreated control cells. From differential expression data and differential degradation, we calculated differential transcriptional synthesis by summing log_2_ (fold-change) of transcript degradation with log_2_ (fold-change) of transcript abundance.

### Functional annotation and GSEA

Transcript functional annotation assigned by matching transcripts to the MSigDB Hallmark Collection^21^. We used the fgsea (v.1.30.0) package to compute enrichment scores across the full collection for differential expression, degradation, and transcription^40^. To score activating transcription factor enrichment, we filtered the DoRothEA (v.1.7.4) collection of human regulons for consensus transcriptional regulators and used the resulting list of gene targets as gene sets for input in GSEA^24^. In all cases, we applied a false discovery rate threshold of 25%. Leading edge genes were extracted from the fgsea results for membership comparison across conditions.

### m^6^A quantification analysis

We computed per-transcript coding region m^6^A modification rates from GLORI datasets deposited by Liu et al.^41^. Using datasets from HeLa cells cultured under both standard conditions and hypoxia, we aggregated m^6^A modification rates across the longest coding isoform of each transcript by summing modification density at each site across the full coding region. We then divided that modification sum by the length of the coding region to obtain a modification density per kilobase. We related these results to our differential decay data by computing Spearman’s correlation coefficient across all transcripts for both normoxia and hypoxia GLORI datasets and in each of our conditions: STM2457, hypoxia, and both together.

### Cross condition concordance of differential expression data

In assessing whether m^6^A and hypoxia have a mechanistic relationship, we plotted differential expression for STM2457 treatment in normoxia against its effects in hypoxia; as well as differential expression for hypoxia in DMSO against the context of STM2457. To assess the concordance of these treatments, we computed Pearson’s correlation for all transcripts in each condition. Additionally, we computed a linear fit between genes with significantly different differential expression in at least one condition. To assess whether transcripts with low concordance across conditions shared a functional relationship, we chose a threshold of 1.4 absolute difference in log_2_ fold-change between conditions and performed over-representation analysis against the KEGG pathway database.

