## Supplemental Figures for "METTL3 inhibition attenuates the macrophage type I interferon response"

### Supplemental Information

**Authors:**

Edward M.C. Courvan, Cody J.S. Hecht, Frederick Longshore-Neate, Luisa M. Vasconcelos, Roy Parker

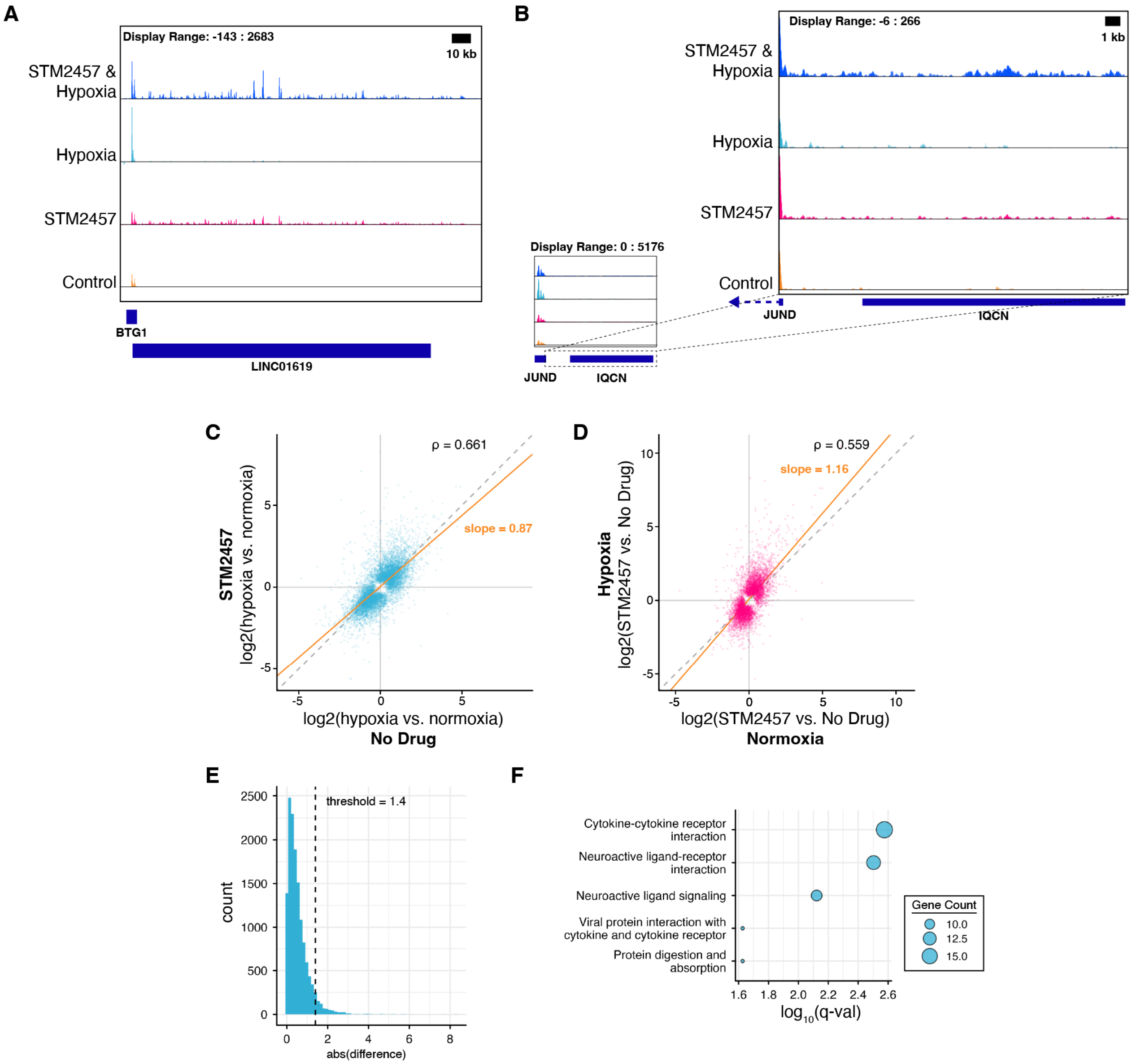


**Figure S1. STM2457 treatment induces transcriptional readthrough.** Genome browser tracks for BTG1 (A) and JUND (B) and their downstream annotations showing how STM2457 leads to mapped reads in lncRNAs and coding genes. C&D) Differential expression for either hypoxia (C) or STM2457(D) treatment plotted in the context of the other. Yellow line indicates a linear fit of transcripts with significantly ($q < 0.05$) different expression in at least one axis. ρ indicates transcriptome wide correlation between contexts. E), Histogram of absolute differences between gene expression changes induced by hypoxia in either STM2457 treated or untreated THP1 macrophages. Threshold of 1.4 was determined by scanning for the maximal sensitivity of over enrichment of KEGG pathway sets. F) Enriched KEGG gene sets among genes with greater than 1.4 absolute difference in expression change as described in E.

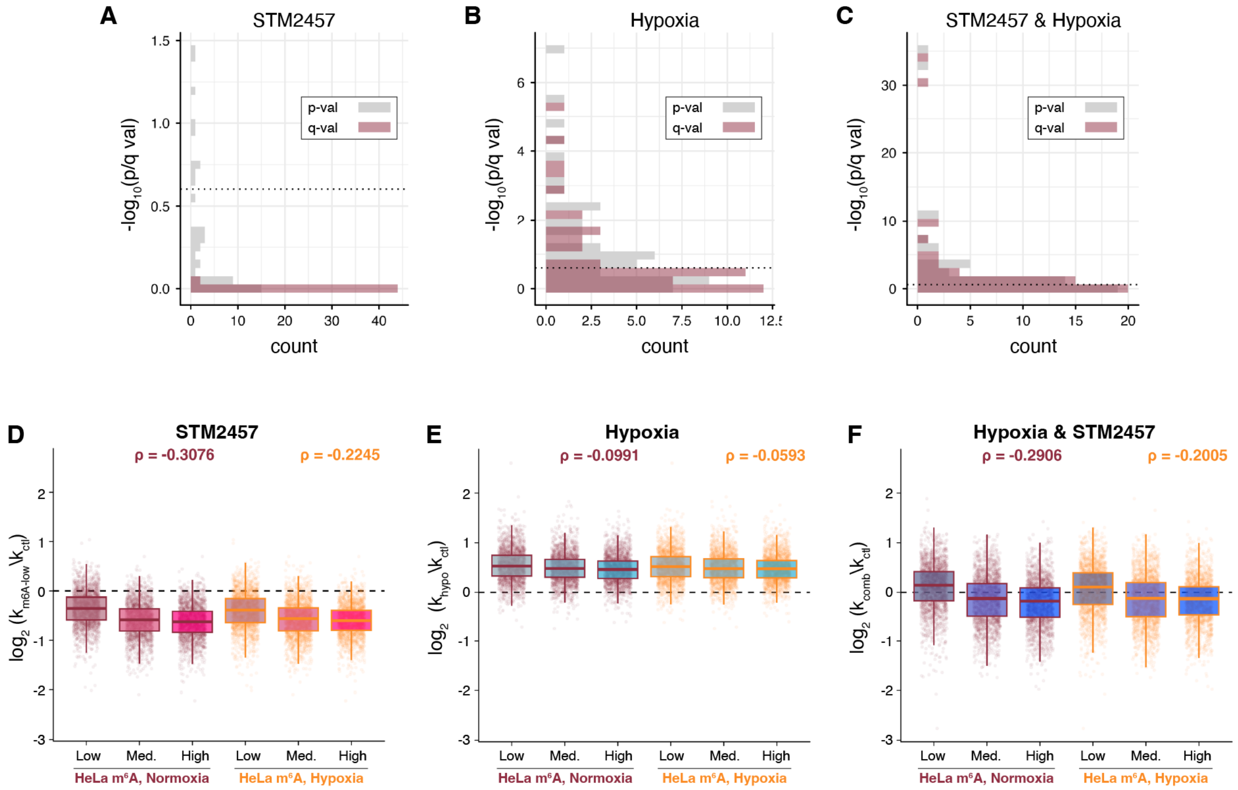


**Figure S2. METTL3 inhibition is pathway agnostic.** A-C) Histograms of raw p-values and false discovery rate adjusted q-values for GSEA on differential degradation analysis of STM2457 (A), Hypoxia (B), or both together (C). Dotted line indicates 25% FDR threshold. D-F) Differential decay rates measured with TimeLapse-seq in D) METTL3 inhibition with STM2457, E) hypoxia, or F) both treatments together plotted against $m^6A$ modification rates taken from GLORI datasets generated from either normoxic or hypoxic HeLa cells. Low, medium (med.) and high $m^6A$ bins were determined by separating transcripts into three quantiles based on their GLORI $m^6A$ levels. ρ reports Spearman's rank order correlation between GLORI $m^6A$ level and differential degradation.

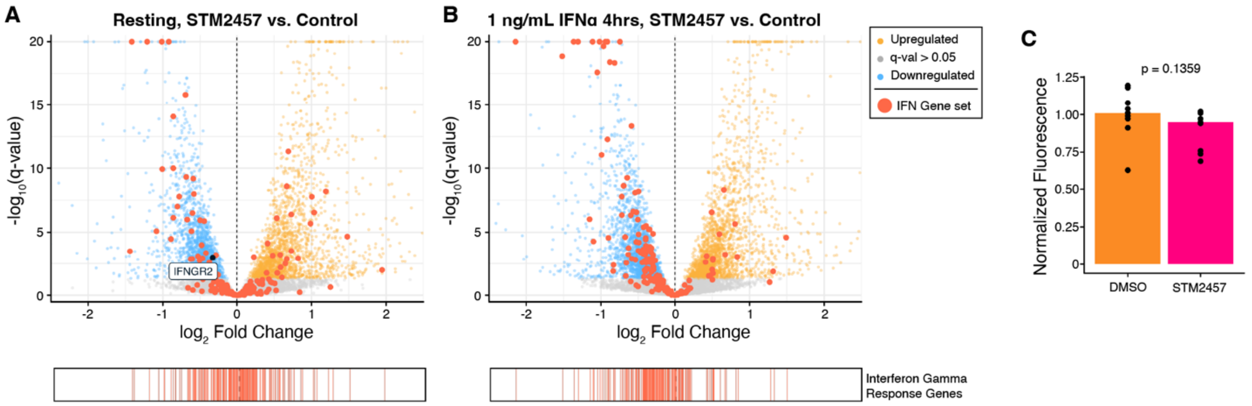


**Figure S3. Interferon-γ related gene expression is muted by STM2457.** A&B) PolyA RNA-seq of resting THP1 macrophages (A) or THP1 macrophages treated with 1 ng/mL interferon-α for four hours (B). Significance determined by an FDR threshold of 25%. Transcripts belonging to the MSigDB hallmark "Interferon Gamma Response" gene set are noted in red and scored in the strip plot below. C) Flow cytometry for interferon gamma receptor 2 in THP1 macrophages in either control or STM2457 treatment. Reported p value taken from Mann-Whitney U. Fluorescence values were background subtracted and normalized to DMSO control.

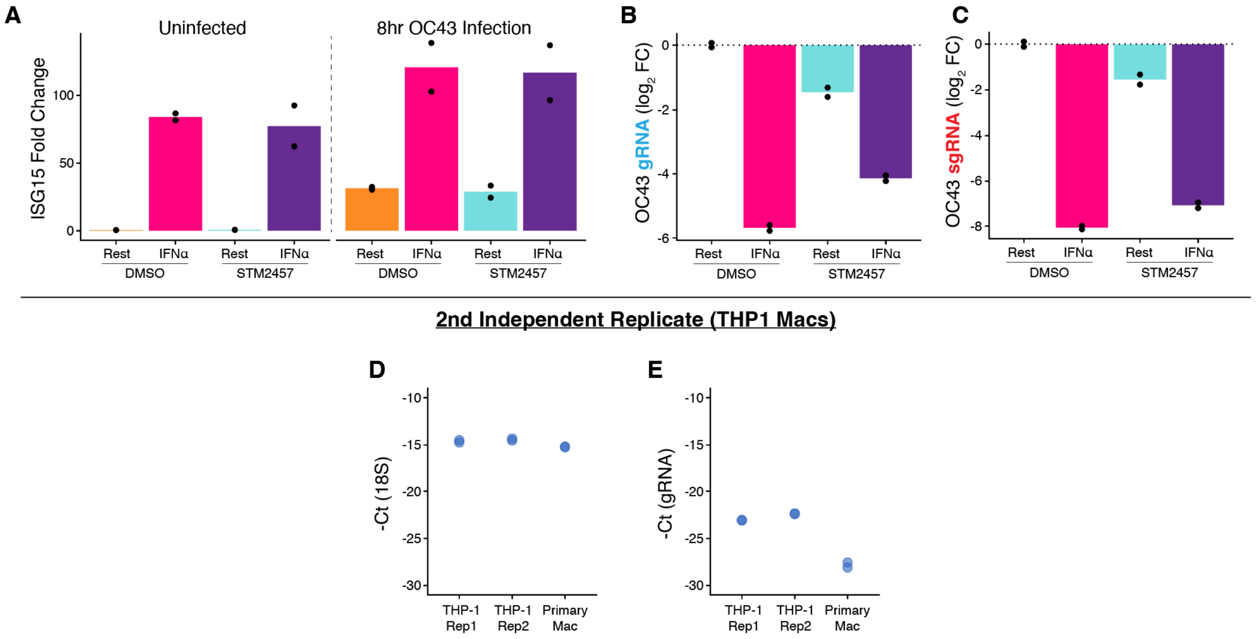


**Figure S4. Macrophages are susceptible to OC43 under METTL3 inhibition.** A) RT-qPCR for ISG15 across all four treatment conditions in uninfected and OC43 infected cells. B) RT-qPCR for OC43 gRNA. C) RT-qPCR for OC43 sgRNA. D&E) Inverted $C_t$ values for 18S probes (D) and gRNA probes (E) in the resting DMSO control condition for all three experiments.

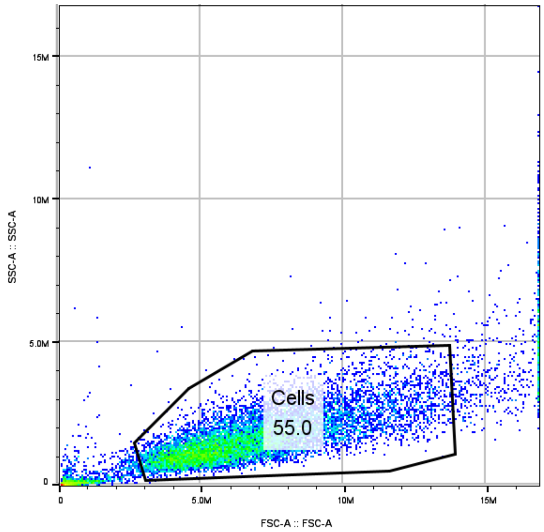


**Figure S5. Gating of THP1 macrophages.** Unlabeled control population of cells used to define the FSC-A/SSC-A Gate used for all samples.

| Primer Set | Forward | Reverse |
|---|---|---|
| ISG15 | CGCAGATCACCCAGAAGATCG | TTCGTCGCATTTGTCCACCA |
| OC43 RdRp | GAGTGTAGATGCCCGTCTCG | ATCAACACGCTGAAAACGGC |
| OC43 NP | TACGGCACCGATATTGACGG | GTGCGCGAAGTAGATCTGGA |
| 18S | CGGCGACGACCCATTCGAAC | GAATCGAACCCTGATTCCCCGTC |

**Table S1. Primer pairs used in this study.** OC43 RdRp is annotated as "gRNA" and OC43 NP is annotated as amplifying the "sgRNA" population.

**Supplemental Data 1.** Summary tables of differential expression, decay, and synthesis data from TimeLapse-seq data presented in this manuscript (Figures 1&2)

**Supplemental Data 2.** Summary tables of differential expression from polyA 3' end seq presented in this manuscript (Figure 3).
